# Micro-Pharmacodynamics: Bridging *In Vitro* and *In Vivo* Experimental Scales in Testing Drug Efficacy and Resistance

**DOI:** 10.1101/031294

**Authors:** Katarzyna A. Rejniak

**Author notes:** K. A. Rejniak is with the Integrated Mathematical Oncology Department of the H. Lee Moffitt Cancer Center & Research Institute, Tampa, FL 33612 & the Department of Oncologic Sciences, College of Medicine of the University of South Florida. Tampa. FL 33612.

## Abstract

Systemic chemotherapy is one of the main anticancer treatments used for most kinds of clinically diagnosed tumors. However, the efficacy of these drugs can be hampered by the physical attributes of the tumor tissue that can impede the transport of therapeutic agents to tumor cells in sufficient quantities. As a result, drugs that work well *in vitro* often fail at clinical trials when confronted with the complexities of interstitial transport within the tumor microenvironment. The *microPD* model that we developed is used to investigate the penetration of drug molecules through the tumor tissue and influenced by the physical and metabolic properties of tumor microenvironment, and how it affects drug efficacy and the emergence of drug resistance.

## I. Introduction to Micro-Pharmacodynamics

The complexity of tumor microenvironment, both physical (interstitial fluid pressure, tissue cellular architecture, fibril composition, or extracellular matrix stiffness) and metabolic (oxygen, glucose, lactate, acidic pH, or growth factors levels) results in various physiological barriers to the transport of drugs or imaging agents into the tumor tissue [1–3]. The classical models of pharmacokinetics and pharmacodynamics (PK/PD) represent tissues and organs as homogeneous well-mixed compartments. However, both normal and tumor tissues are heterogeneous in their structure and response to metabolites and treatments. We present here a biomechanical model (called *microPD*; the microenvironment-influenced pharmaco-dynamics model) of the interstitial transport of both metabolites and drugs, and their role in tumor growth and its response to treatments.

## II. Illustrative Results of MicroPD Applications

The *microPD* model has been initially developed to test the role of tumor tissue architecture, which is defined by its porosity (the amount of void space between the cells) and cellular density (the number of cells per a given volume), in the interstitial transport of therapeutic or imaging agents [4]. We have shown that the typical measure of transport phenomena, the Peclet number (a ratio of the advective to diffusive transport, where advection is due to the fluid flow, and diffusion is driven by a drug gradient), is not indicative of the dominant transport mode for the moderate Peclet values. In these cases, the interstitial transport can be either advection- or diffusion-dominated depending on the tumor tissue architecture; that is, whether it is composed of small or large cells. Here, we discuss the applications of the *microPD* model to projects that were part of either the ICBP or PSOC grants, or have bridged both projects scopes.

### A. Enhancing the efficacy of HAPs (PSOC)

We used the *microPD* model, after its calibration to the properties of the MiaPaCa-2 pancreatic cancer cells, to formulate a hypothesis about how the exogenous pyruvate (a metabolic modifier shown to increase oxygen uptake in cells) could enhance the efficacy of the TH-302 hypoxia-activated pro-drug (HAP) [5]. We showed that within a patch of tissue with the normoxia–hypoxia border (N-HB) stabilized at a distance of 110 μm from the vasculature, the bolus injection of both pyruvate and TH-302 leads to the decrease in tissue oxygenation by 30% (the N-HB shifts left to a distance of 76 μm) in 10 min after injection. As a result, the region of the inter-tumoral hypoxia, and thus the region of activation of the HAPs is increased three-fold resulting in an elevated cell death (88% increase when compared to TH-302 alone) in 30 min after injection (Figure 1). This model is now being used to design optimal schedules of combined therapies that utilize three or more therapeutic compounds. As the behavior of HAPs is complex, it is beyond intuition how to determine the right dosage and the right order of drugs applied in combination to maximize the efficacy of TH-302. Our model provides a means to examine multiple scheduling options before testing them in a laboratory.

**Figure 1.**
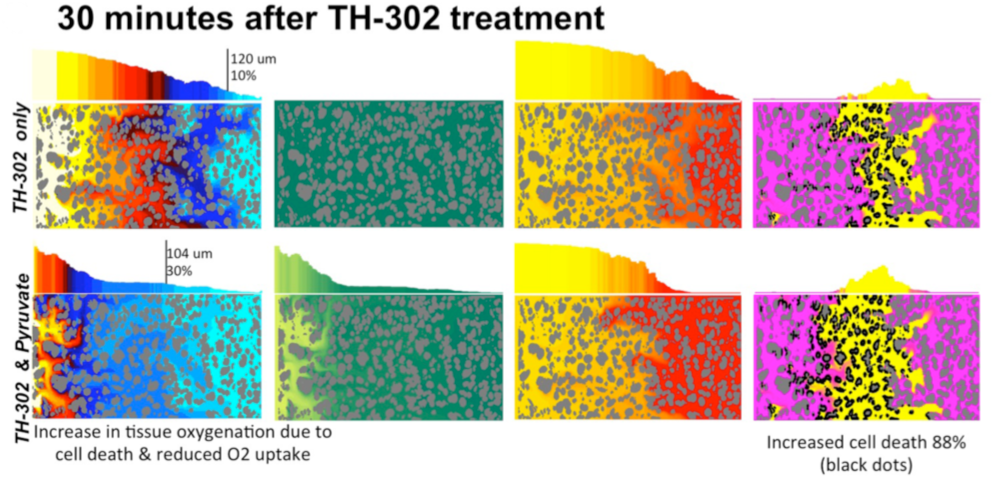
Oxygen and active TH-302 distributions in simulated tumors calibrated to MiaPaCa-2 tissue histology. *Adapted from [5].*

### B. Predicting the effects of cell-cycle targeted drugs (ICBP)

A version of the *microPD* model has been used to develop an *in silico* analogue of the 2D cell colony culture technique. The model is quantitatively calibrated to fit data from lung adenocarcinoma PC9 cells treated with different levels of *erlotinib* [6]. By exploring the model parameter space, we proposed three hypotheses on how cell migratory, quiescent and proliferative properties change if cells are exposed to different drug concentrations (Figure 2). However, in *in vitro* experiments, all cells in a single well are typically exposed to a uniform concentration of the drug, and the cell response to different drug concentrations is measured by varying drug levels among different wells. By contrast, tumor cells growing within the tissue can be exposed to concentrations that vary both in time and in space.

**Figure 2.**
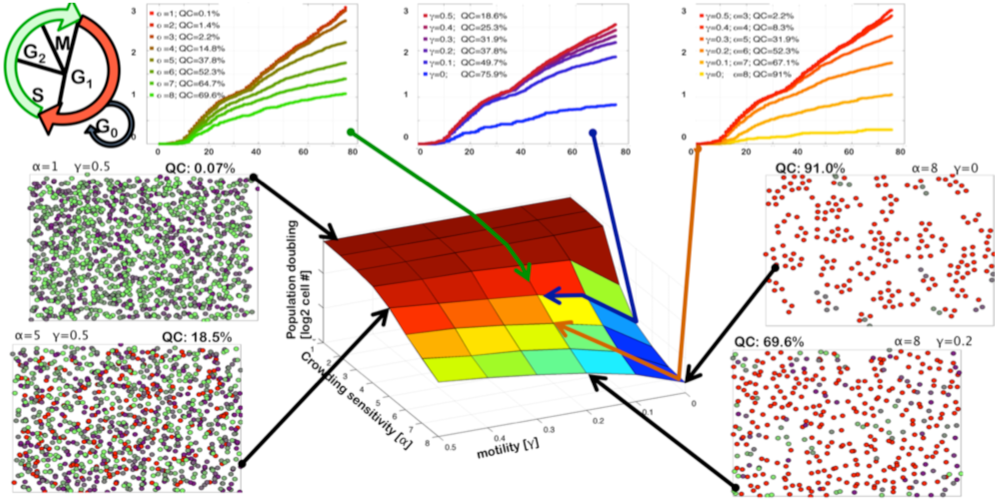
A parameter space of the *microPD-cell culture* model with three hypothetical cases of cell motility and quiescence modified in response to varied drug concentrations. *Adapted from [7].*

To bring *in vitro* data to an *in vivo* context we applied the *microPD-tissue scale* model in which drug is supplied from a non-uniform vasculature leading to a drug-gradient build-up over time. Therefore, in a single computer simulation, tumor cells are exposed to different drug levels depending on their location, and a single cell can be exposed to varied drug levels at different time points. Our results show that for the three proposed scenarios of cell phenotypic evolution, the final configurations of *in vivo*-like simulations differ in both cell age distributions and cell cycle phase distributions [7]. An extension of this model has been applied to investigate how the efficacy of cell-cycle checkpoint inhibitors is affected by the formation of tight tumor clusters versus sparse tumor colonies both in 2D and 3D cultures [8].

### C. Microenvironmental niches & drug resistance (WhAM!)

The approaches developed in the two previous projects were applied to investigate how complex gradients of metabolites and drugs, which result from a chaotic tumor vasculature, form specific microenvironmental niches that promote the emergence of anti-cancer drug resistance [9]. Our simulations show (Figure 3) that the spatial regions characterized by either low-drug/sufficient oxygen (low D) or low-drug/low-oxygen (H & low D) levels have a significant impact on the transient and long-term tumor behavior when drug resistance occurs as a result of drug action (acquired resistance).

**Figure 3.**
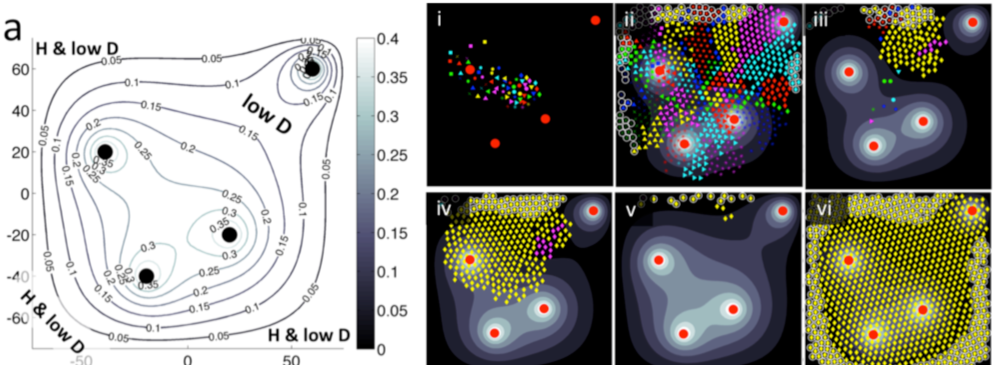
(a) Microenvironmental niches of low drug level (low D) or low drug & hypoxia level (H & low D) resulting in spatial evolution of a population of drug resistant tumor cells (i-vi). *Adapted from [9].*

## III. Quick Guide to the Mathematical Methods

The *microPD* model uses a combination of the classic fluid–structure interaction method of the regularized Stokeslets [10], the lattice-free agent-based model of individual cells [9], and the diffusion–advection–reaction equations for the kinetics of all metabolites and drugs.

In this model, we explicitly consider the cellular and vascular architecture of the tumor tissue, the interstitial fluid flow, and the concentrations of both metabolites and drug molecules.

### A. Equations

The Stokes equations (Figure 4a,b) are used due to small Reynolds numbers associated with the physical problems under investigation. The regularized forces applied to the vasculature and cell boundaries create the physiologically relevant interstitial fluid flow. The interactions between individual cells are defined using linear repulsive Hookean springs (Figure 4c), and cell motion is governed by a Newtonian equation (Figure 4d). The kinetics of all metabolites and drugs includes the diffusive and advective motions, the cellular uptake and the transition from an inactive to activated state (Figure 4e-g).

**Figure 4.**
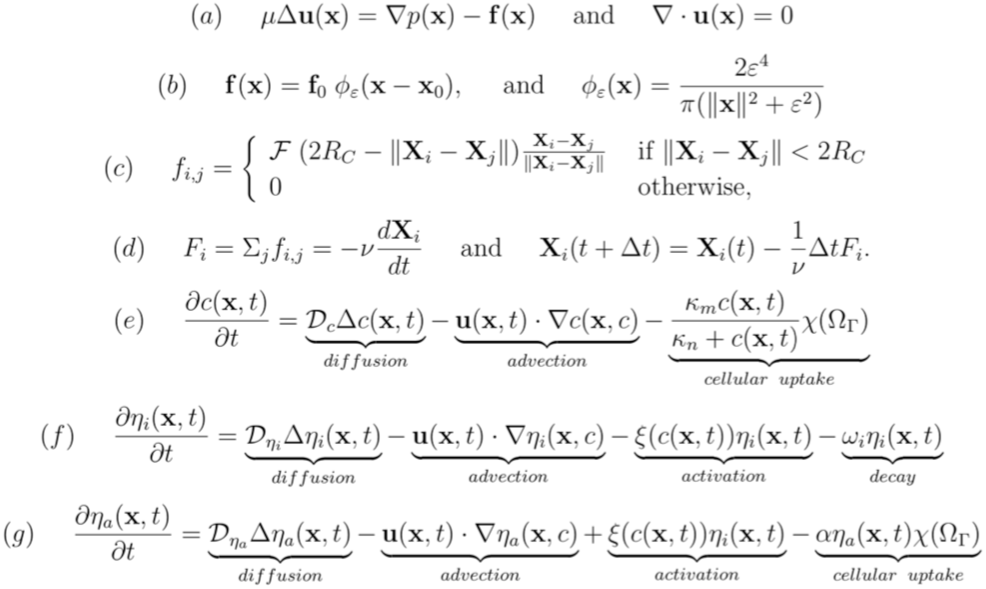
Model equations: (a-b) regularized Stokeslets; (c) Hookean repulsive springs; (d) Newtonian cell motion; (e-f) diffusion-advection-reaction kinetics. *Adapted from [5] and [9].*

### B. Type of settings in which these methods are useful

The presented model enables investigations of the behavior of individual cells, and their interactions with one another and with the surrounding microenvironment. Both the mechanical and chemical clues sensed by the cells can be included in the model, as well as multiple cellular types and the many metabolite concentrations. Many intracellular properties, including the cell cycle or heterogeneous cell sensitivity to the surrounding metabolic landscape and drug concentrations can be also incorporated in the model. The *microPD* model can be used to represent the *in vitro* cell cultures and the *in vivo* tumor tissues, and is an ideal computational tool to bridge the different (*in vivo, in vitro,* or *ex vitro*) experimental scales.

## Acknowledgment

K.A.R would like to thank her experimental collaborators Drs. V. Quaranta and D. Tyson from Vanderbilt ICBP, Drs. R. Gillies and J. Wojtkowiak from Moffitt PSOC, and mathematical modelers, members of the WhAM! project Drs. J. Gevertz from the College of New Jersey, J. Perez-Velazquez from Helmholtz Zentrum Munich, K.A. Norton from Johns Hopkins University, A. Volkening from Brown University, and Z. Aminzare from Rutgers University.

## Notes

Research supported by the NIH Physical Sciences-Oncology Centers Grant LT54-CA-143970, and the NIH Integrated Cancer Biology Program Grant U54-CA-113007.

